# The priming phosphorylation of KaiC is activated by the release of its autokinase autoinhibition

**DOI:** 10.1101/2024.03.21.584037

**Authors:** Yoshihiko Furuike, Yasuhiro Onoue, Shinji Saito, Toshifumi Mori, Shuji Akiyama

## Abstract

KaiC, a cyanobacterial circadian clock protein with autokinase activity, catalyzes the dual phosphorylation of its own S431 and T432 residues in a circadian manner in the presence of KaiA and KaiB. Priming phosphorylation at T432 is a key step that promotes secondary phosphorylation at S431. Although KaiA binding is considered essential for KaiC phosphorylation, the mechanisms underlying the activation and inactivation of priming phosphorylation remain elusive. We found that the priming phosphorylation proceeds even in the absence of KaiA, but is autoinhibited within KaiC, which decreases the rate constant to 0.019 h^-1^. The autoinhibition of KaiC and the mechanism underlying the release from autoinhibition by KaiA were examined by KaiC structural analysis, and by classical molecular dynamics and quantum mechanics / molecular mechanics simulations. We found that the side chain of T432 adopts two rotamers in dephosphorylated KaiC, one of which places T432 in a position suitable for a nucleophilic attack on the terminal phosphate of adenosine triphosphate (ATP). However, the nucleophilicity of T432 was insufficient to overcome an energy barrier of approximately 22 kcal mol^-1^ because the catalytic function of a nearby base, E318, was self-suppressed by hydrogen bonding to positively charged R385. Biochemical assays of KaiC mutants showed that the autoinhibition of KaiC autokinase activity is attenuated by conferring T432 high nucleophilicity through the KaiA-assisted release of R385 from E318 to E352. During the circadian cycle, R385 switches interacting partners to inactivate/activate the autokinase function and to ensure the unidirectionality of the KaiC phosphorylation cycle.

**Significance Statement:** KaiC, a central player in the circadian clock system of cyanobacteria, undergoes an ordered phosphorylation cycle in the presence of KaiA and KaiB. To elucidate the mechanism underlying the rhythmic regulation of the KaiC autokinase, we performed structural analyses, computational simulations, and biochemical assays of KaiC and its mutants. The results indicate that KaiC is essentially an autoinhibited autokinase, and the autoinhibition of primary phosphorylation at its T432 residue is attenuated by conferring it high nucleophilicity against the terminal phosphate of adenosine triphosphate. KaiA contributes to releasing the autoinhibition of KaiC in a morning phase by switching the interacting partners of R385 from a catalytic glutamate E318 to E352, as well as ensuring unidirectionality of the KaiC phosphorylation cycle.

## Introduction

Circadian clocks are time-keeping systems that enable organisms to adapt to daily changes in environmental conditions by modifying intracellular processes. The clock systems from bacteria to mammals share three physiological characteristics (1): self-sustained oscillation with an approximately 24 h period, an endogenous period of temperature insensitivity (temperature compensation), and a phase entrainment in response to external stimuli. These properties depend on complex and diverse biochemical events involving clock proteins and clock-related components (2, 3).

Phosphoryl modification is one of the main post-translational events regulating clock proteins. Phosphorylation of the clock proteins modulates their function, atomic-scale structure, and dynamics, thereby controlling the stoichiometry and stability of multimeric clock-protein complexes (4, 5). Rhythmic phosphorylation of the clock proteins is observed in many organisms (6, 7) as well as *in vitro* reconstituted systems (8, 9). To achieve a circadian rhythm of phosphorylation, at least two processes for activating and inhibiting phosphorylation must be present, and these processes must switch appropriately from phase to phase.

Protein kinases phosphorylate targeted amino acids such as Ser, Thr, and Tyr in a substrate molecule or in the kinase itself (autokinase). Kinases can be constitutively active or switch between the active and inactive forms through intermolecular interactions or autophosphorylation (10). The activity of constitutively activate kinases with external substrates can be modulated by the concentration of the substrate. However, the activity of autokinases is controlled by modifying the active site structure or the intramolecular accessibility to the targeted amino acids.

KaiC is a component of the cyanobacterial circadian clock of *Synechococcus elongatus* PCC 7942 that possesses autokinase activity. KaiC is composed of tandemly duplicated N-terminal (CI) and C-terminal (CII) domains (11), and it forms a double-ring hexamer by incorporating one molecule of ATP or ADP at every CI–CI and CII–CII interface (12). In the presence of KaiA and KaiB, the S431 and T432 residues in the CII domain of KaiC are phosphorylated/dephosphorylated in a circadian manner (13). The protein undergoes a phosphorylation cycle starting in a fully dephosphorylated form (KaiC-ST) and changing to a T432-phosphorylated form (KaiC-SpT: priming phosphorylation), a S431/T432-phosphorylated form (KaiC-pSpT: secondary phosphorylation), and a S431-phosphorylated form (KaiC-pST) before returning to the KaiC-ST form (14, 15). Studies indicate that the phosphoryl modification of KaiC-ST proceeds only after the addition of KaiA (13). KaiA does not show amino acid sequence or structure similarity with other existing kinases, suggesting that binding of KaiA to KaiC is necessary for the autophosphorylation of KaiC. Although the function and structure of the KaiA–KaiC complex have been studied extensively (16–18), the dynamic nature of the KaiA–KaiC complex has made these studies difficult, and the mechanism underlying the regulation of KaiC autokinase activity remains to be elucidated.

In the absence of KaiA, KaiC is present in the KaiC-ST form under physiological conditions (19–21); therefore, the priming phosphorylation of T432 is frequently discussed in relation to the KaiA binding (13, 18). However, in this study we found that KaiC is autophosphorylated in the absence of KaiA, albeit at a slower rate. Although this finding is not consistent with the conventional interpretation that KaiC cannot be auto-phosphorylated in the absence of KaiA, it suggests that the priming phosphorylation of KaiC is activated by releasing the autoinhibition of autophosphorylation activity through interaction with KaiA. This finding provides an opportunity to examine the mechanism by which KaiA activates KaiC autophosphorylation by analyzing the autoinhibition of KaiC without using KaiA.

In this study, we revisited our previous crystal structure of KaiC-ST to examine the mechanism underlying the autoinhibition of KaiC autophosphorylation. The priming phosphorylation of KaiC at T432 was examined using classical molecular dynamics (MD) and quantum mechanics/molecular mechanics (QM/MM) simulations, and the energy barrier for the phosphoryl transfer reaction mediated by the identified catalytic base was estimated. Amino acid substitutions in the catalytic base and regulatory residues nearby suggest a mechanism by which the KaiC autophosphorylation is autoinhibited, and this inhibition is released by interaction with KaiA.

## Results

### Priming Phosphorylation at T432 in KaiC Occurs Even in the Absence of KaiA, Albeit at a Slower Rate

At a standard experimental temperature of 30°C (19–21), KaiC underwent the autodephosphorylation and was eventually detected in a fully dephosphorylated from, KaiC-ST (*Fig. 1A* and *SI Appendix*, Fig. S1). After 40 h of incubation at 30°C, a small fraction of KaiC remained in the KaiC-SpT, KaiC-pSpT, and KaiC-pST forms, which did not change detectably after another 20 h.

**Figure 1.**
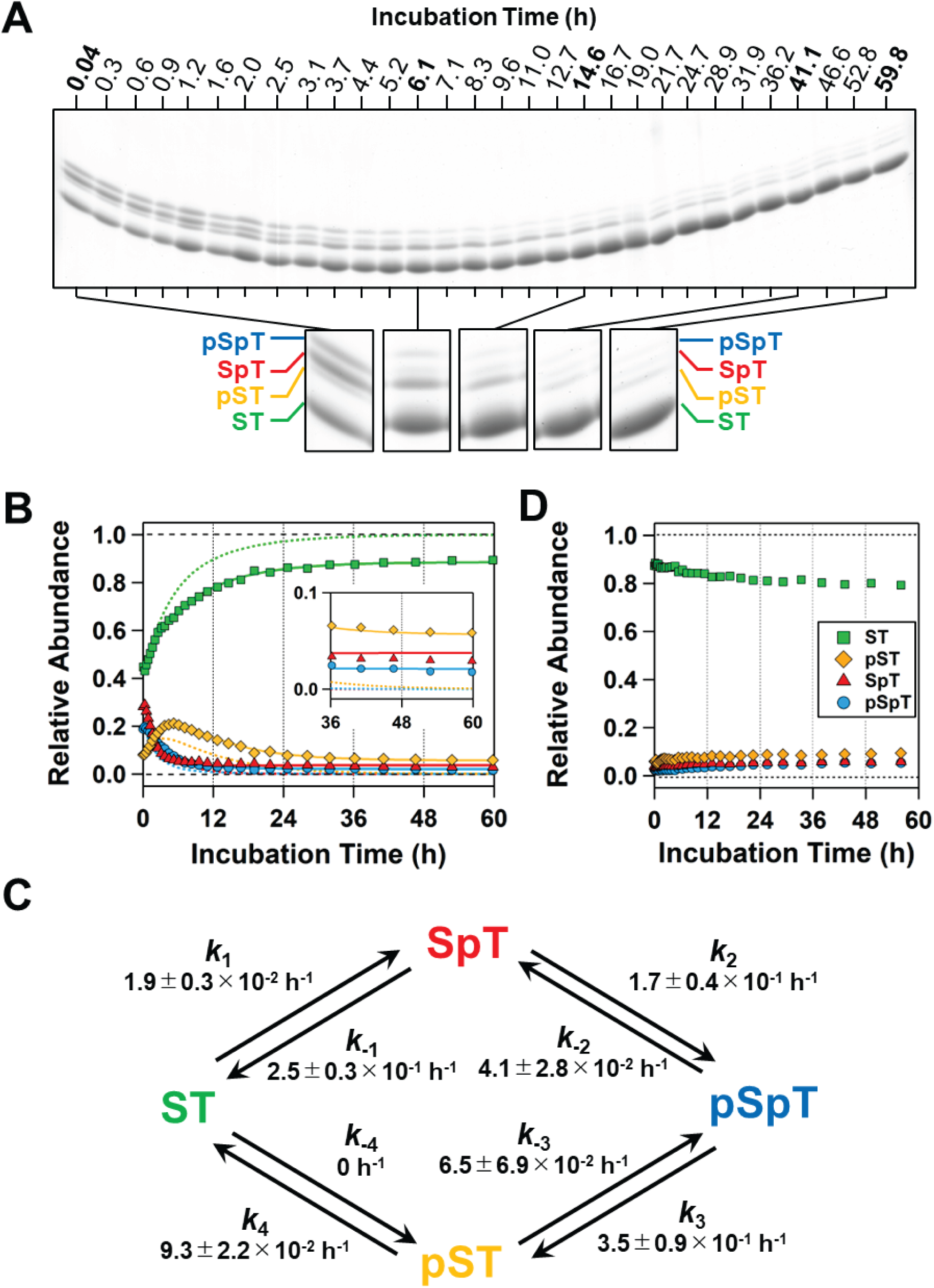
Priming phosphorylation of T432 in KaiC in the absence of KaiA. (*A*) Representative image of an SDS-PAGE gel loaded with samples from KaiC under autodephosphorylation conditions at 30°C. To improve the visualization of the four bands corresponding to ST, SpT, pSpT, and SpT, the lanes containing the samples collected at 0.04, 6.1, 14.6, 41.1, and 59.8 h are shown in enlarged images. The whole image of the SDS-PAGE gel is available in *SI Appendix*, Fig. S1. (*B*) Densitometric analysis of the gel image in panel *A* was performed to determine the time course of the relative abundance of the ST (green squares), SpT (red triangles), pSpT (blue circles), and SpT (orange diamonds) forms. The solid lines represent the best fit from global nonlinear least-squares analysis using the four-state model in panel *C*. The inset shows the enlarged plots corresponding to the longer period region. Converged rate constants are compiled in *SI Appendix*, Table S1. The dashed lines represent the worse fit obtained by forcing the rate constants for priming (*k*_1_: ST → SpT) and secondary (*k*_2_: SpT → pSpT) phosphorylation to zero and leaving the others unchanged. (*C*) Reversible four-state cyclic model for KaiC phosphorylation/dephosphorylation. Each step (x = 1, 2, 3, and 4) consists of forward (*k*_X_) and backward (*k*_-X_) reactions. (*D*) Autophosphorylation of KaiC was induced at 30°C in the absence of KaiA by increasing the pH from 8.0 to 9.0. The original data for this plot are shown in *SI Appendix*, Fig. S3.

Next, we performed a densitometric analysis of the gel image (Fig. 1*A*) and carefully examined the resultant kinetic traces for autodephosphorylation (Fig. 1*B*). Consistent with previous reports (19–21), the KaiC-SpT and KaiC-pSpT fractions decreased rapidly, whereas the KaiC-pST fraction increased transiently and then began to decrease, and the KaiC-ST form increased progressively. At 40–60 h of incubation, the fractions of KaiC-SpT, KaiC-pSpT, and KaiC-pST remained constant at approximately 0.03, 0.02, and 0.06, respectively (Fig. 1*B*, Inset). KaiA was previously reported to be essential for the autophosphorylation of KaiC-ST at 30°C (13). However, the present observations imply that KaiC-ST is autophosphorylated even in the absence of KaiA, albeit at a slower rate.

As shown by solid lines in Fig. 1*B*, the global-fitting of a reversible four-state cyclic model (14, 15) was used to examine the autodephosphorylation dynamics of KaiC, and the eight associated rate constants were determined (Fig. 1*C*). The contributions of the priming (KaiC-ST → KaiC-SpT, *k*_1_ = 1.9 ± 0.3 × 10^-2^ h^-1^) and secondary (KaiC-SpT → KaiC-pSpT, *k*_2_ = 1.7 ± 0.4 × 10^-1^ h^-1^) autophosphorylation events were considerable and reproducible (n = 5, *SI Appendix*, Table S1), even when the reaction was initiated using different relative amounts of KaiC in the four states (*SI Appendix*, Fig. S2). When both *k*_1_ and *k*_2_ were forced to zero while keeping the others unchanged, the fitting to the residual phosphorylated species became considerably poorer (Fig. 1*B*, dashed lines). This result means that the small but non-zero values of *k*_1_ and *k*_2_ make a significant contribution to the autodephosphorylation dynamics in Fig. 1*B*. Consistent with this interpretation, KaiC, which was dephosphorylated by equilibration at pH 8 for 50 h at 30°C, was slightly but obviously phosphorylated after the pH was increased to 9.0 (Fig. 1*D* and *SI Appendix*, Fig. S3).

The present data and analysis suggest that the priming phosphorylation of KaiC-ST occurs slowly but continuously in the absence of KaiA, even under steady-state conditions in which KaiC-ST is the predominant form over other phosphorylated forms at 30°C (Fig. 1*B*). As illustrated in the model in Fig. 1*C*, approximately 60% [*k*_-1_ / (*k*_-1_ + *k*_2_)] of the minor KaiC-SpT form is directly converted back to KaiC-ST; however, the remaining 40% undergoes secondary autophosphorylation and is restored to the KaiC-ST form by autodephosphorylation in a cyclic manner. At 30°C, a small amount of KaiC-SpT, the product of the priming phosphorylation, undergoes a repeated cycle of formation and loss even in the absence of KaiA.

### One of Two Rotamers of T432 in KaiC-ST is Near Ready for the Autokinase Substrate

The molecular mechanism of the priming phosphorylation remains under debate partly due to the limited spatial resolution of the KaiA–KaiC complex (16–18). The slow but steady priming phosphorylation of KaiC in the absence of KaiA suggests that the structure of KaiC-ST itself could be used to examine this mechanism.

We thus inspected the existing crystal structure of KaiC-ST (22) from the perspective of priming phosphorylation. An ATP molecule bound between two neighboring CII protomers (CII-ATP, Fig. 2*A*) serves as the phosphate source for the kinase reaction. S431, T432, and a phospho-switch (PSw) belonging to one protomer, together with E318*, E352*, and R385* belonging to a second protomer (hereafter, marked with asterisks), constitute an active site around CII-ATP (Fig. 2*B*). The PSw located at the upstream of S431 undergoes a helix-coil transition according to the phosphoryl modification essentially at S431 (22). In KaiC-ST, the helical conformation of the PSw facilitates positioning of T432 closer to CII-ATP than S431 (Fig. 2*B*), which is favorable for the priming phosphorylation reaction (22).

**Figure 2.**
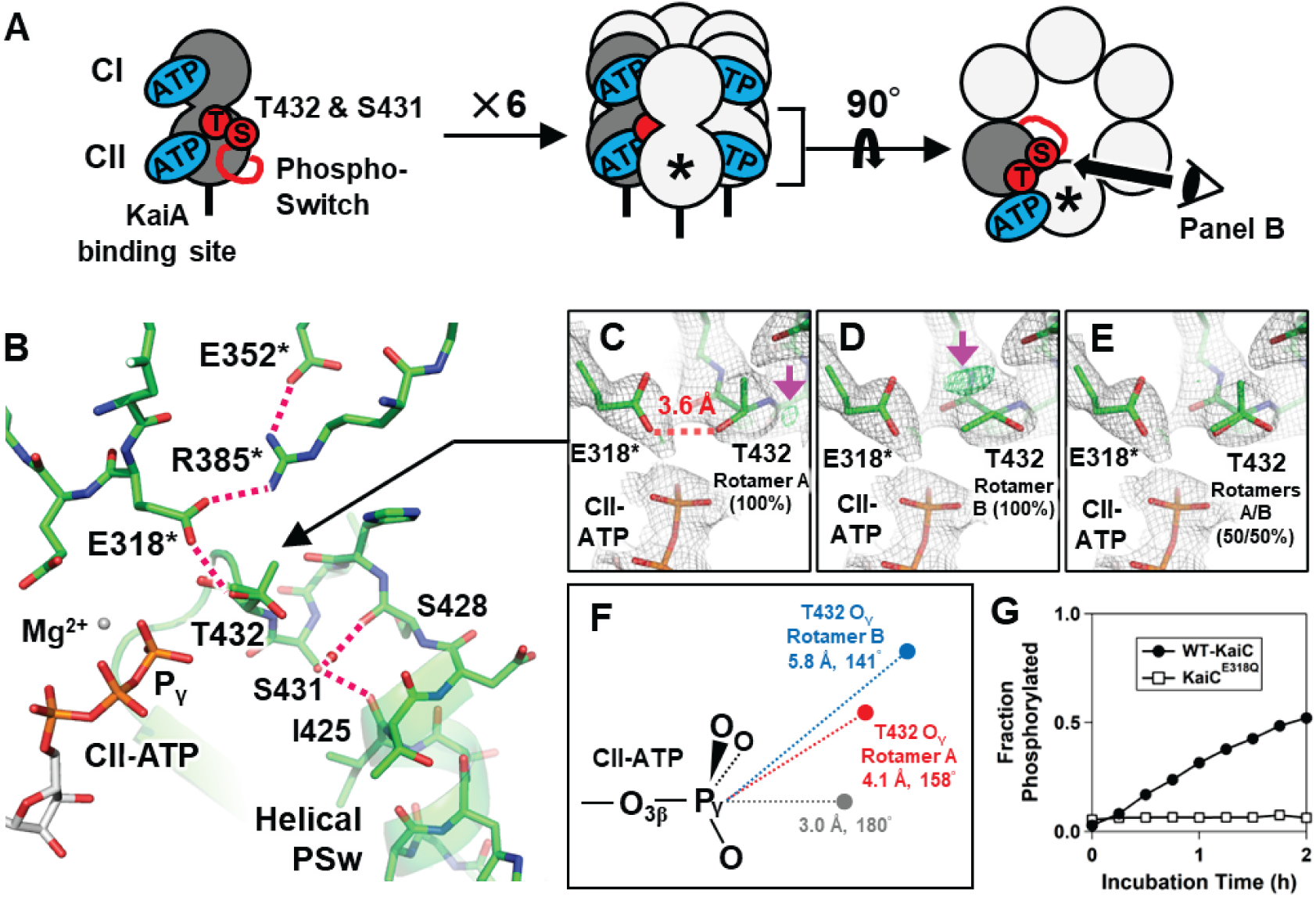
Structure of KaiC-ST. (*A*) Overall structure of the KaiC-ST hexamer. (*B*) Enlarged image of the autokinase active site around CII-ATP. Hydrogen bonds between the phosphorylation sites (S431 and T432), E318*, R385*, E352*, and PSw are indicated by magenta dashed lines. Residues from a neighboring protomer are indicated by asterisks. (*C*–*E*) Occupancy of two rotamers of the T432 side chain. The 2|Fo|–|Fc| (gray mesh) and |Fo|–|Fc| difference maps (green mesh) are countered at 1.0- and 3.0-sigma levels, respectively. Magenta arrows indicate the emerged peaks in the |Fo|–|Fc| difference maps. (*F*) Schematic drawing of the positions of T432 O_γ_ in rotamers A and B. (*G*) Time courses of phosphorylated fraction for wild-type KaiC and KaiC^E318Q^ after KaiA addition (0.04 mg/mL) at time zero.

The side chain of T432 adopted two conformations corresponding to a rotation of 120° around the C_α_ – C_β_ axis. Refinement using only one rotamer (rotamer A), in which the T432 O_γ_ was oriented toward the phosphorus atom (P_γ_) of the γ-phosphate group of CII-ATP, yielded a small but obvious positive difference Fourier peak near T432 C_β_ (magenta arrow in Fig. 2*C*). Refinement using the other rotamer (rotamer B) showed a similar but separate peak with the T432 O_γ_ directed away from CII-ATP (Fig. 2*D*). The occupancy of each rotamer was estimated at approximately 50% (Fig. 2*E*).

T432 O_γ_ of rotamer A (T432^rot-A^ O_γ_) was nearly configured as a suitable substrate for kinase reactions. In general, Thr or Ser reacts with an ATP molecule through an S_N_2-like transition structure (Fig. 2*F*), in which the nucleophilic O_γ_ is brought as close as possible to the ATP P_γ_ along with the ATP P_γ_ – O_3β_ axis to form an O_γ_ – ATP P_γ_ bond, and the ATP P_γ_ – O_3β_ bond is then cleaved in a concerted manner. T432^rot-A^ O_γ_ was only 4.1 Å away from the ATP P_γ_ with a T432^rot-A^ O_γ_ –CII-ATP P_γ_ – CII-ATP O_3β_ angle of 158° (Fig. 2*F*), which is an almost optimal in-line position (∼3 Å and 180°) for the reaction. T432^rot-B^ O_γ_ was in a less favorable position (5.8 Å and 141°) than T432^rot-A^ O_γ_.

To become a ready-to-react nucleophile, the hydroxyl group of T432^rot-A^ has to be deprotonated by a general base. In KaiC-ST, T432^rot-A^ O_γ_ could potentially be hydrogen-bonded to E318* O_ε_ with a distance of 3.6 Å (Fig. 2*C*). In fact, the E318Q substitution inhibited the priming phosphorylation in the presence of KaiA (Fig. 2*G*), suggesting that E318* O_ε_ attracts T432^rot-A^ H_γ_ through a hydrogen-bonding interaction, and T432^rot-A^ O_γ_ becomes the reactive nucleophile. These structural analyses suggest that the rotamer A is the structural basis for the slow but steady priming phosphorylation of KaiC-ST.

### QM/MM and MD Simulations of a Proton Transfer and T432 Phosphorylation Reaction

To visualize the priming phosphorylation independent of KaiA, we performed QM/MM metadynamics (metaD) simulation using the KaiC-ST hexamer with T432^rot-A^ as the initial coordinate. In the simulations, we used a QM region consisting of the side chains of T432^rot-A^, E318*, R385*, and part of CII-ATP/Mg^2+^ (Fig. 3*A*, represented by green or orange stick models).

**Figure 3.**
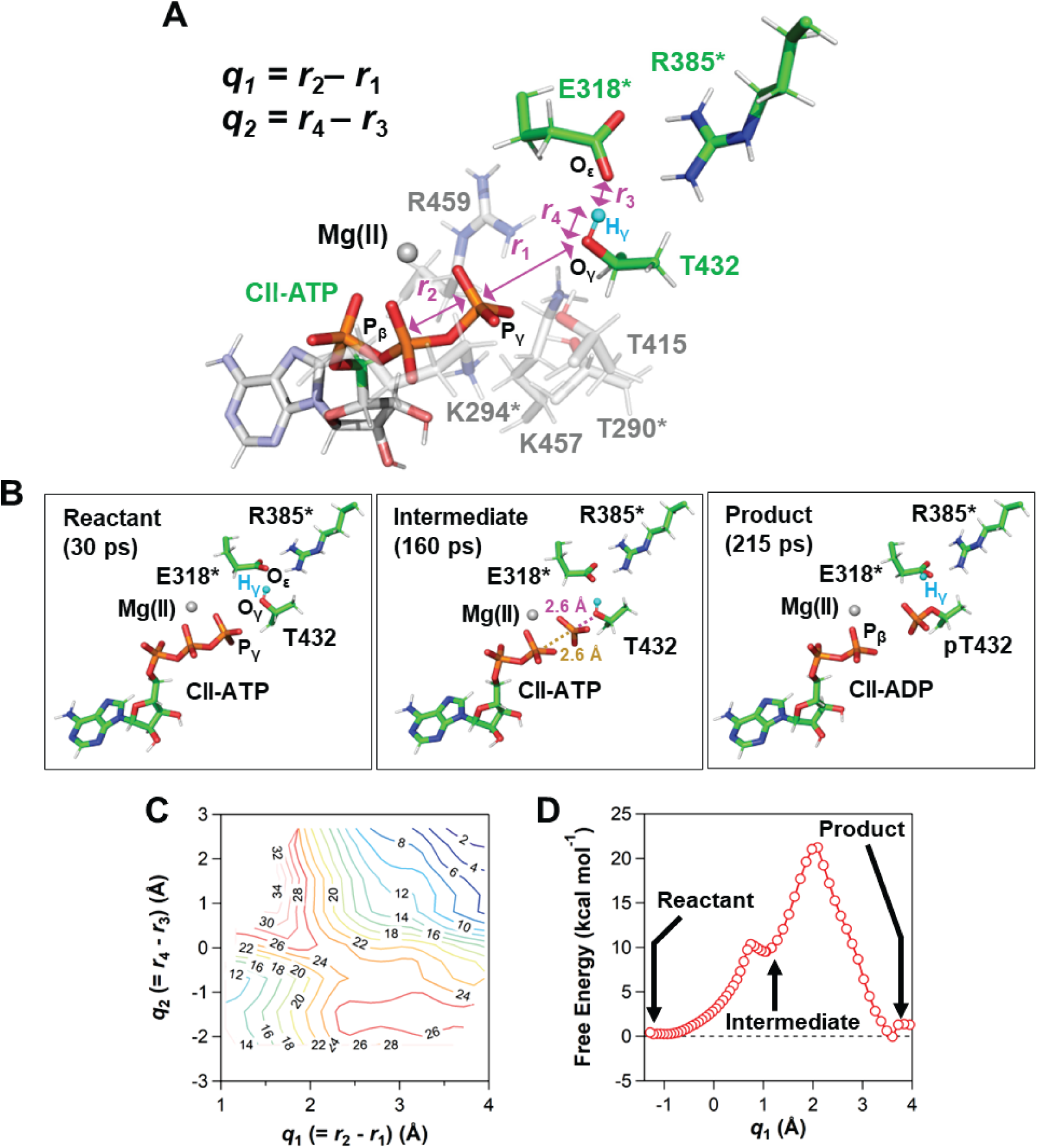
QM/MM simulations of the phosphoryl transfer from CII-ATP to T432. (*A*) Definition of the QM region and reaction coordinates. The QM region consists of the side chains of T432, E318*, and R385*, triphosphate and part of the ribose group (C5’, H5’1, and H5’2) of CII-ATP, and Mg^2+^ (green or orange stick models). The hydrogen atom (T432 H_γ_) which is transferred from T432 to E318* is highlighted (a cyan sphere). Reaction coordinates are defined as *q*_1_ = *r*_2_ – *r*_1_ (*r*_1_: CII-ATP P_γ_ – T432 O_γ_, *r*_2_: CII-ATP P_γ_ – CII-ATP P_β_) and *q*_2_ = *r*_4_ – *r*_3_ (*r*_3_: T432 H_γ_ – E318* O_ε_, *r*_4_: T432 H_γ_ – T432 O_γ_). (*B*) Structures obtained during the QM/MM metaD simulation of T432 phosphorylation in KaiC-ST. (*C*) Two-dimensional free energy surface calculated using the QM region defined in panel *A*. (*D*) Free energy surface along *q*_1_ from the QM/MM calculation.

The QM/MM metaD suggested that T432^rot-A^ O_γ_ is activated as a nucleophile and attacks CII-ATP P_γ_ in synergy with E318*. As shown in Fig. 3*B*, E318* O_ε_ attracted the configured T432^rot-A^ H_γ_ through a hydrogen-bonding interaction prior to dissociating the γ-phosphate from CII-ATP, thereby directing the hydroxyl of T432^rot-A^ to the ready-to-react orientation (reactant in Fig. 3*B*). The obtained trajectory shows that the CII-ATP P_γ_ – CII-ATP O_3β_ bond dissociated to form an intermediate state at 160 ps. After remaining in the intermediate state, the system moved to a transition state by decreasing the T432^rot-A^ O_γ_ – E318* O_ε_ distance to ∼3 Å; at this moment, a covalent bond was formed between P_γ_ and T432^rot-A^ O_γ_ concomitant with the transfer of T432^rot-A^ H_γ_ to E318* (product in Fig. 3*B*). This suggests that both the formation of the P_γ_ – T432^rot-A^ O_γ_ bond and the proton transfer from T432^rot-A^ O_γ_ to E318* O_ε_ must be considered when describing the transition state of the phosphoryl transfer step.

To identify possible reaction pathways, we calculated the two-dimensional free energy surface as a function of the relative positions of CII-ATP P_γ_ and T432^rot-A^ H_γ_. As the reaction proceeded from the intermediate state (lower left in Fig. 3*C*), the CII-ATP P_γ_ – T432^rot-A^ O_γ_ distance (*r*_1_) became shorter, whereas the CII-ATP P_γ_ – CII-ATP P_β_ distance (*r*_2_) increased (*r*_2_ – *r*_1_ = *q*_1_ increased). After the *q*_1_ value reached 2.5, the attraction of T432^rot-A^ H_γ_ by E318* O_ε_ shortened the T432^rot-A^ H_γ_ – E318* O_ε_ distance (*r*_3_) but lengthened the T432^rot-A^ H_γ_ – T432^rot-A^ O_γ_ distance (*r*_4_) (*r*_4_ – *r*_3_ = *q*_2_ increased). The most plausible reaction pathways emerged as diagonal clefts from the lower left to the upper right corners in Fig. 3*C*. Together with the one-dimensional free energy surface connecting the reactant and intermediate states (Fig. 3*D*), the present simulations suggest a barrier as high as approximately 22 kcal mol^-1^ to yield KaiC-SpT from KaiC-ST.

In a separate MD simulation, the side chain of T432 in five of the six protomers of KaiC-ST was re-oriented to fit the rotamer B during the simulation, whereas one survived as rotamer A for 1 µs mostly within approximately 5 Å from CII-ATP P_γ_ (*SI Appendix*, Fig. S4). This implies that the ready-to-react structures with the short T432^rot-A^ O_γ_ – CII-ATP P_γ_ distance exist in solution in a short-lived state, i.e., a conformationally excited state (23). The high energy barrier and the limited population of ready-to-react structures suggest that the priming phosphorylation of T432 in KaiC-ST is not efficient in the absence of KaiA. This is consistent with experimental observations that only a small amount of pT432 is produced from KaiC-ST at 30°C (Fig. 1*A* and 1*B*). The accumulation of KaiC-SpT in the presence of KaiA (Fig. 2*G*) suggests that KaiA plays a role in lowering the energy barrier by enhancing the nucleophilic character of T432^rot-A^ O_γ_.

### R385*–E352* Hydrogen Bonding Pair Indirectly Regulates the Nucleophilicity of T432^rot-A^ O_γ_ for Priming Phosphorylation

The nucleophilicity of T432^rot-A^ O_γ_ can be enhanced by strongly withdrawing its proton, T432^rot-A^ H_γ_. The negatively charged carboxylate moiety of E318* appears to be in an ideal position for this purpose. However, its negative charge is partially neutralized by hydrogen bonding with the positively charged guanidino moiety of R385* that is further stabilized by negatively charged E352* (Fig. 2*B*). We studied the range of the indirect effects on T432^rot-A^ O_γ_ by examining the hydrogen-bonding network in KaiC-ST from T432^rot-A^ O_γ_ to E318*, R385*, and even E352* (Fig. 2*B*).

The role of R385* itself was previously discussed based on the observations (24) that the R385A mutant (KaiC^R385A^) accumulated as mainly phosphorylated states *in vivo* but as the dephosphorylated state without KaiA *in vitro*. We revisited the effect of the R385A mutation by analyzing the autodephosphorylation dynamics of KaiC^R385A^ with the global-fitting analysis (Fig. 4*A*). The effect of harboring the guanidino moiety at the 385th position was most obvious in the process of the priming phosphorylation (*t*-test, *p* = 1.1 × 10^-5^) (*SI Appendix*, Fig. S5), as the R385A mutation doubled the *k*_1_ value even without KaiA (*k*_1_ = 3.8 ± 0.2 × 10^-2^ h^-1^) (Fig. 4*B*). This result suggests that R385* neutralizes the negative charge of E318*, eventually suppressing the nucleophilicity of T432^rot-A^ O_γ_ in KaiC-ST. At the same time, the priming phosphorylation activity of KaiC^R385A^-ST was further accelerated in the presence of KaiA (*k*_1_ = ∼2.0 × 10^-1^ h^-1^) (Fig. 4*C* and 4*D*), although it was not as high as the activity of KaiA-stimulated KaiC-ST (*k*_1_ = ∼3.5 × 10^-1^ h^-1^) (Fig. 4*E*). A ratio of the *k*_1_ value in the presence of KaiA to the *k*_1_ value in the absence of KaiA was 18 for KaiC-ST, while it considerably decreased to be approximately 5 for KaiC^R385A^-ST (Fig. 4*C*). The release of R385* from E318* is necessary to trigger the priming phosphorylation, but not sufficient to increase the nucleophilic reactivity of T432^rot-A^ up to the KaiA-stimulated state of KaiC-ST.

**Figure 4.**
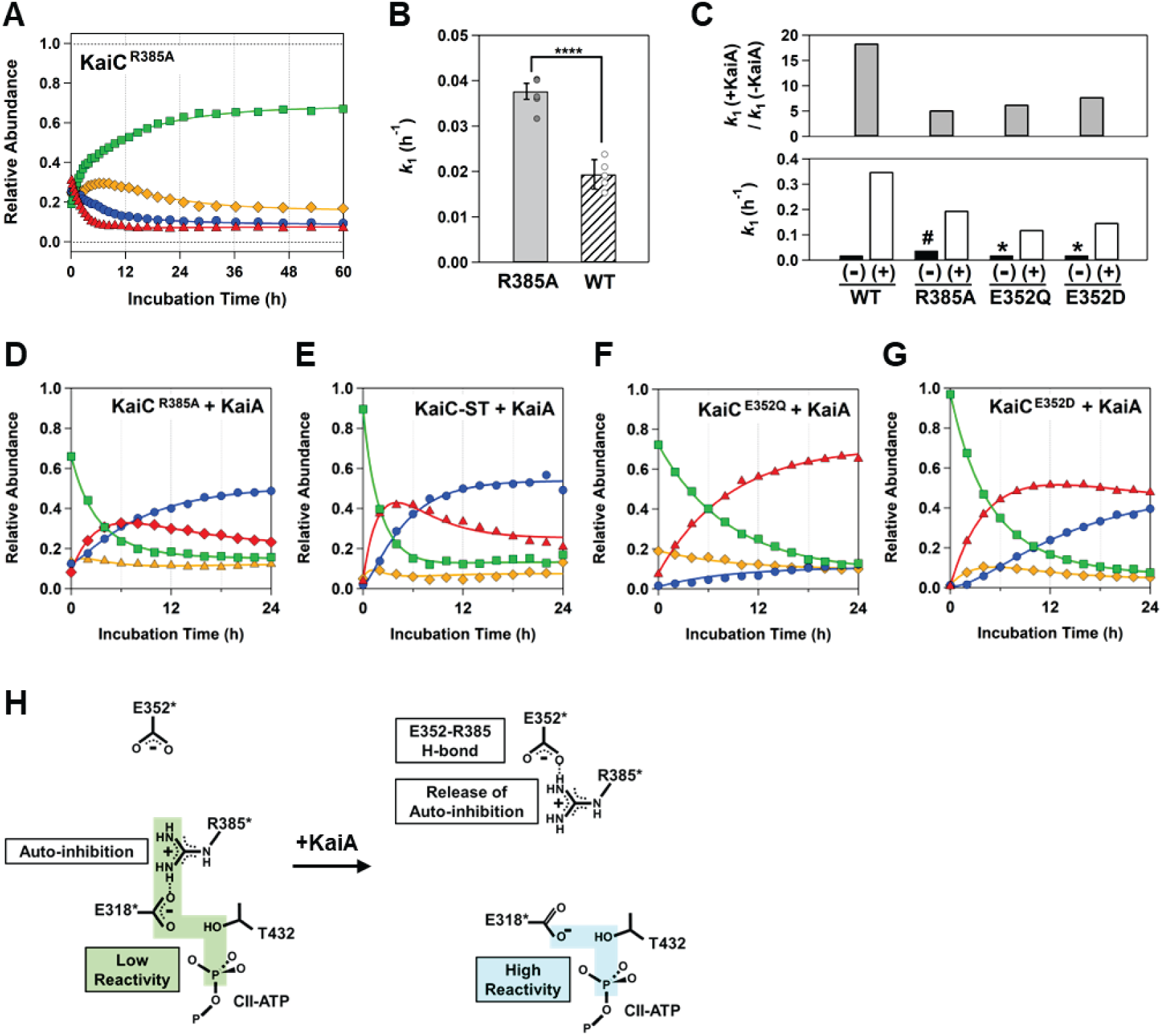
Local and non-local effects on the nucleophilicity of T432^rot-A^ O_γ_. (*A*) Autodephosphorylation dynamics of KaiC^R385A^ (ST: green squares, SpT: red triangles, pST: orange diamonds, and pSpT: blue circles). The solid lines represent the best fit from global nonlinear least-squares analysis using the four-state model in Fig. 1C. (*B*) Effect of the R385A substitution on the rate (*k*_1_) of the priming phosphorylation. *t* test; ****more significant (P < 0.0005). See details in *SI Appendix*, Fig. S5. (*C*) Ratios of the *k*_1_ value in the presence of KaiA (+) to the *k*_1_ value in the absence of KaiA (-). ^#^0.038 h^-1^. *The same *k*_1_ value as wild-type KaiC-ST (0.019 h^-1^) is assumed for KaiC^E352Q^ and KaiC^E352D^. (*D*–*G*) Time courses of four phosphorylation states of KaiC^R385A^ (+KaiA), KaiC-ST (+KaiA), KaiC^E352Q^ (+KaiA), and KaiC^E352D^ (+KaiA), respectively, at 30°C. The correspondence of the fitted lines and graph markers is the same as in panel *A*. KaiC-ST, KaiC^E352Q^, KaiC^E352D^, and KaiC^R385A^ (0.2 mg/mL) were pre-incubated at 30°C for 48 h and then at time zero mixed with KaiA at a final concentration of 0.04 mg/mL. (*H*) Schematic drawing of possible hydrogen bonds (black dotted line) and T432^rot-A^ H_γ_-withdrawing reactivity of E318* to enhance the nucleophilicity of T432^rot-A^ O_γ_.

We found that the hydrogen-bond formation R385* and E352* indirectly contributes to the activation by KaiA. A KaiC^E352Q^ mutant in which the charge at the 352nd position was neutralized showed a markedly reduced rate of decrease in the ST state in the presence of KaiA (*k*_1_ = ∼1.2 × 10^-1^ h^-1^) (Fig. 4*F*) compared with wild-type KaiC (Fig. 4*E*). Similar observations were made for a KaiC^E352D^ mutant (*k*_1_ = ∼1.5 × 10^-1^ h^-1^) (Fig. 4*G*) in which the side chain at the 352nd position remained negatively charged, but not sufficiently long enough to form a hydrogen bond with R385*. One possible common reason for the reduced *k*_1_(+KaiA) / *k*_1_(-KaiA) ratios of KaiC^R385A^, KaiC^E352Q^, and KaiC^E352D^ (Fig. 4*C*) is the destabilization of the KaiA-induced state with R385* captured by E352* (Fig. 4*H*). In the crystal structure of a S431A/T432A mutant (KaiC-AA), which is currently interpreted as the state mimicking KaiC-ST bound by KaiA (25), the side chain of R385* is separated from E318* by a distance of 5.2 Å (Fig. 5*B*) and trapped by E352* through the hydrogen bonds (Fig. 5*A*). We speculate that high nucleophilicity T432^rot-A^ O_γ_ is continuously achieved by at least two different types of the KaiA-assisted conformational changes of KaiC-ST; locally liberating R385* from E318* and non-locally forming the hydrogen bonds between R385* and E352*.

**Figure 5.**
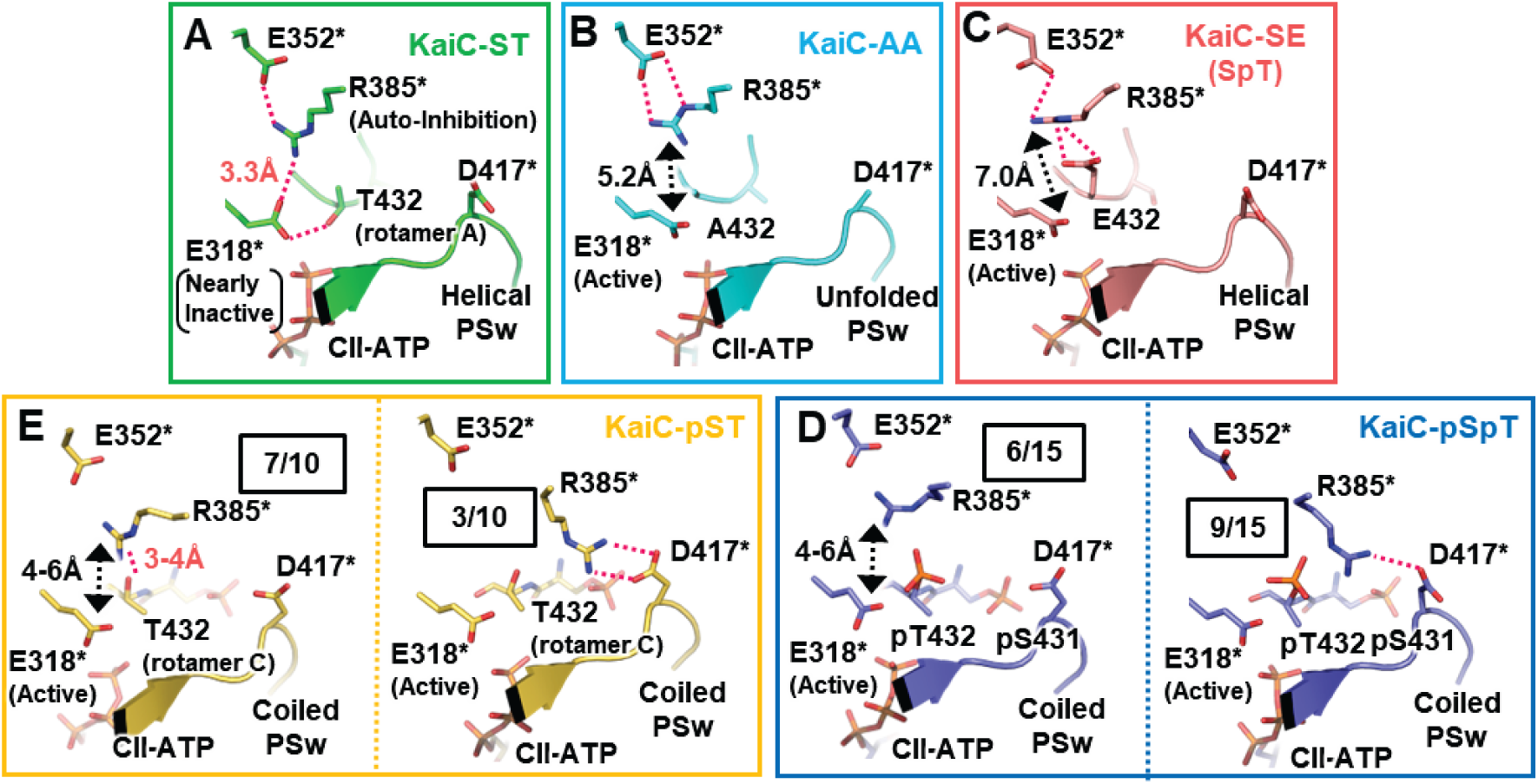
R385* rearrangement during the circadian cycle. The side chain orientations of R385* in (*A*) KaiC-ST (PDB code: 7DYJ), (*B*) KaiC-AA (PDB code: 7DYE), (*C*) KaiC-SpT (PDB code: 7DYI), (*D*) KaiC-pSpT (PDB code: 7DXQ, 7DY1, and 7V3X), and (*E*) KaiC-pST (PDB code: 7DY1, 7V3X, and 7DY2) are shown. The numbers enclosed in boxes indicate the fractions of the protomers with the corresponding conformation of R385* in the structure library. Hydrogen bonds are indicated by magenta dashed lines. Inter-atomic distances beyond hydrogen bonding distance are indicated by black dashed arrows.

### R385* Rearrangement Assures the Unidirectionality of the KaiC Phosphorylation Cycle

The existing library of the KaiC structures (22, 25) suggests the crucial role of R385* rearrangement even after priming phosphorylation. In the T432E mutant (KaiC-SE), which is considered as the best KaiC-SpT mimetic (14, 26), R385* is pulled away from E318* by hydrogen bonds with E432 and E352* (Fig. 5*C*), leaving E318* activated as the reactive base suitable for attracting a hydrogen atom. This orientation of R385* in KaiC-SE may be effective in promoting the E318*-mediated activation of S431 O_γ_ for the secondary phosphorylation reaction.

In nine of the fifteen pSpT protomers in the library (Fig. 5*D*, right), the positively charged guanidino moiety of R385* was trapped by the negatively charged side chain of D417*. Although R385* in the other six protomers remained within the interaction range of E352*, the distance between R385* and E318* was considerably longer than the hydrogen bonding distance (Fig. 5*D*, left). These findings suggest that E318* of KaiC-pSpT remains as the active free base, despite the lack of reactive side chains for the kinase reaction around CII-ATP. In such an arrangement of R385*, however, KaiC-pST could be autophosphorylated again and re-converted to KaiC-pSpT. In seven of the ten pST protomers in the library, the side chain of T432 in KaiC-pST, which may restrain this reverse reaction, formed a third rotamer C (Fig. 5*E*) with its hydroxyl group oriented away from E318* and trapped by R385* (Fig. 5*E*, left).

The coil-to-helix transition of PSw (22), which is promoted by the autodephosphorylation of pS431, returns R385* to the configuration observed in KaiC-ST (Fig. 5*A*). During the circadian cycle, the positively charged guanidino group of R385* swaps interacting partners among E318*, E352*, and D417* to achieve inactivation/activation of T432^rot-A^ O_γ_, thereby ensuring the unidirectionality of the phosphorylation cycle of KaiC.

## Discussion

Although KaiA is considered essential for the phosphorylation of KaiC (13), the present findings clearly demonstrate that the priming phosphorylation of T432 occurs at 30°C in the absence of KaiA, albeit at a slower rate. Even under conditions that promote the steady-state predominance of KaiC-ST, <10% of KaiC is transiently expressed as partly or fully phosphorylated forms and constantly cycling among the four states (Fig. 1*C*). The low abundance of these forms is mainly due to inefficient priming phosphorylation, which is limited by the three factors described below.

The first is the marginal stability of the helical PSw in KaiC-ST in solution. The structure of the PSw changed from a helical to a non-helical form in five of six protomers in the QM/MM simulation over a period of 1 µs. This observation is also consistent with our recent mutation study (27), which identified structural polymorphisms of the PSw in KaiC-ST mimetics. The helix-to-coil transition of the PSw causes the hydroxyl group of T432 to face the opposite direction to CII-ATP (Fig. 5*E*), and the resultant location of T432 downstream of the coiled PSw is unfavorable for the kinase reaction.

The second limiting factor is the rotamers of T432 located downstream of the helical PSw. In KaiC-ST carrying the helical PSw, the side chain of T432 shuttles between the two rotamers A and B, waiting for a chance to attack CII-ATP P_γ_. Considering that one half of the rotamers in one-sixth of the KaiC-ST protomers are enabled to react, the fraction of T432 that can undergo the kinase reaction is estimated to be approximately 10%.

This ∼10% of the population is limited by the third factor, an energy barrier that can be as high as 22 kcal mol^-1^. The barrier-crossing mechanism can be largely categorized into base-assisted and substrate-assisted mechanisms (28), which differ in the pathway for withdrawing a proton from the hydroxyl group of Ser or Thr. In the base-assisted mechanism, the substrate proton is transferred to a basic residue nearby, for example from T432^rot-A^ O_γ_ to E318* O_ε_ in KaiC-ST in the absence of KaiA (Fig. 3*B*). By contrast, in the substrate-assisted mechanism, the substrate proton is directly transferred to one of the three oxygen atoms of the γ-phosphate. The following observations support that the priming phosphorylation occurs in a base-assisted manner. The lone pair of E318* is directly oriented toward T432^rot-A^ O_γ_, whereas CII-ATP O_γ_ closest to T432 is already involved in a hydrogen bond with T290* and an electrostatic interaction with K294* (*SI Appendix*, Fig. S6). The current QM/MM metaD simulation also found the hydrogen bond between E318* O_ε_ and T432^rot-A^ H_γ_ prior to the dissociation of the CII-ATP P_γ_ – CII-ATP O_3β_ bond (Fig. 3*B*), and the free energy surfaces further support this mechanism (Fig. 3*D*). Furthermore, the E318Q substitution suppressed the kinase activity in the presence of KaiA (Fig. 2*G*). These factors led us to conclude that the priming phosphorylation of KaiC-ST follows the base-assisted mechanism, regardless of the presence of KaiA.

It is important to discuss our observations with respect to previous QM/MM studies (see (29) and references are therein). QM/MM simulations have been used to investigate phosphoryl transfer reactions in several kinases, including cAMP-dependent protein kinase (30–34), cyclin-dependent kinase (35–37), and N-acetyl-L-glutamate kinase (38, 39). In these examples, the activation energy for the phosphoryl transfer reaction ranges from 10 to 25 kcal mol^-1^. The present QM/MM simulation is the first example of the visualization of atomic-scale details before and after the phosphoryl transfer reaction in clock proteins. The reason for this is that high-resolution structures of fully dephosphorylated clock proteins were not available for a long time, as is the case for KaiC (22). The activation energy for T432^rot-A^ in KaiC-ST, which is approximately 22 kcal mol^-1^, is within the above range, suggesting that once formed, it can be phosphorylated on a reasonable time scale even in the absence of KaiA. According to the Eyring equation, the present experimental rate constant (*k*_1_ = 1.9 ± 0.3 × 10^-2^ h^-1^) for the priming phosphorylation of T432 without KaiA corresponds to an activation energy of 25 kcal mol^-1^. The discrepancy may be due to the setup or the limited accuracy of our QM/MM calculations, although it may also mean that the contribution of the second factor is larger than the above estimate.

The present results suggest that the binding of KaiA attenuates at least one of the limiting factors for the priming phosphorylation of KaiC-ST. The structural and QM/MM evaluations clearly indicate that the carboxylate side chain of E318* attracts T432^rot-A^ H_γ_ and enhances the nucleophilicity of T432^rot-A^ O_γ_ (Fig. 3*B*). This function of E318* as the inherent base is, however, self-suppressed in KaiC-ST by the neutralization of its negative charge with the positively charged guanidino group of R385* nearby. As demonstrated using a series of KaiC mutants (Fig. 4 and Fig. 5*B*), binding of KaiA attenuates this self-suppression by sequestering R385* from E318* to E352*. The observed decay of the KaiC-ST form in the presence of KaiA (Fig. 4*E*) suggests that the rate constant of the priming phosphorylation reaction should increase by a factor of 18 (Fig. 4*C*). This is equivalent to a reduction of the energy barrier of approximately 2 kcal mol^-1^ caused by enhancing the nucleophilicity of the substrate through the KaiA–KaiC interaction. The present results show that the priming autophosphorylation is self-suppressed, although not completely, in KaiC-ST, whereas that KaiA attenuates this self-suppression to achieve the KaiC autophosphorylation on a reasonable time scale as a circadian clock system.

Common protein kinases are maintained mostly inactive in the absence of activating upstream stimuli, whereas inhibitory factors are removed at the biologically appropriate times and locations. As reviewed in detail by Reinhardt and Leonard (10), there are many systems for maintaining protein kinases inactive, such as blocking ATP binding (40), sequestration of substrates or blocking their access (41–44), occupation or occlusion of catalytic cleft (45–47), and binding of regulatory proteins (48).

In most of these examples, positively charged side chains located within hydrogen-bonding distance to the catalytic glutamate/aspartate are extremely limited. Conserved downstream Lys residues that are intrinsically close to the catalytic aspartate (3.3-4.7 Å) are unlikely to form stable hydrogen bonds, regardless of their activation or inactivation, because the carboxylate plane of the catalytic aspartate is almost perpendicular to the orientation of the Lys (*SI Appendix*, Fig. S7*A*). This is also the case for casein kinase 1 delta (CK1d), which is essential for the pacemaker function of mammalian clock systems (49). Insulin receptor kinase (40) and the Src family of tyrosine kinases (43) are exceptions because their catalytic glutamate forms a hydrogen bond with the Arg residue located two or four residues downstream; however, these kinases are not in the ATP-bound form (*SI Appendix*, Fig. S7*B*). This means that, in contrast to KaiC, most protein kinases switch between the active and inactive forms by regulating the substrate access and by autophosphorylating their activation loop (50–53); their catalytic glutamates thus remain unsuppressed to efficiently withdraw a proton from the substrate. KaiC encapsulates both a braking function to fulfill the circadian timescale through the interaction of R385* and E318* and an accelerating function to promote the priming phosphorylation in a morning phase through the interaction of R385* and E352*.

The priming phosphorylation in KaiC can be summarized as follows. The side chain of T432, which was sequestered as the rotamer C in KaiC-pST (Fig. 6*A*), shuttles between two rotamers, A and B, in KaiC-ST to optimize its attack of CII-ATP P_γ_ (Fig. 6*B*). The primary phosphorylation proceeds, albeit slowly, in the ST state even in the absence of KaiA, but its efficiency is low because the function of E318* as the catalytic base to enhance the nucleophilicity of T432 O_γ_ is autoinhibited by R385* (Fig. 6*C*). KaiA binds to the C-terminus of KaiC (16, 17) and releases this autoinhibition by switching the interacting partner of R385* from E318* to E352* (Fig. 6*D*). The increased negative charge of E318* facilitates the attraction of T432^rot-A^ H_γ_, resulting in the concerted proton transfer from T432^rot-A^ to E318* and the nucleophilic attack against CII-ATP P_γ_ by T432^rot-A^ O_γ_ (Fig. 6*D*). In KaiC-SpT (Fig. 6*E*), R385* remains evacuated from E318* by the phosphorylated T432 to promote the secondary phosphorylation at S431.

**Figure 6.**
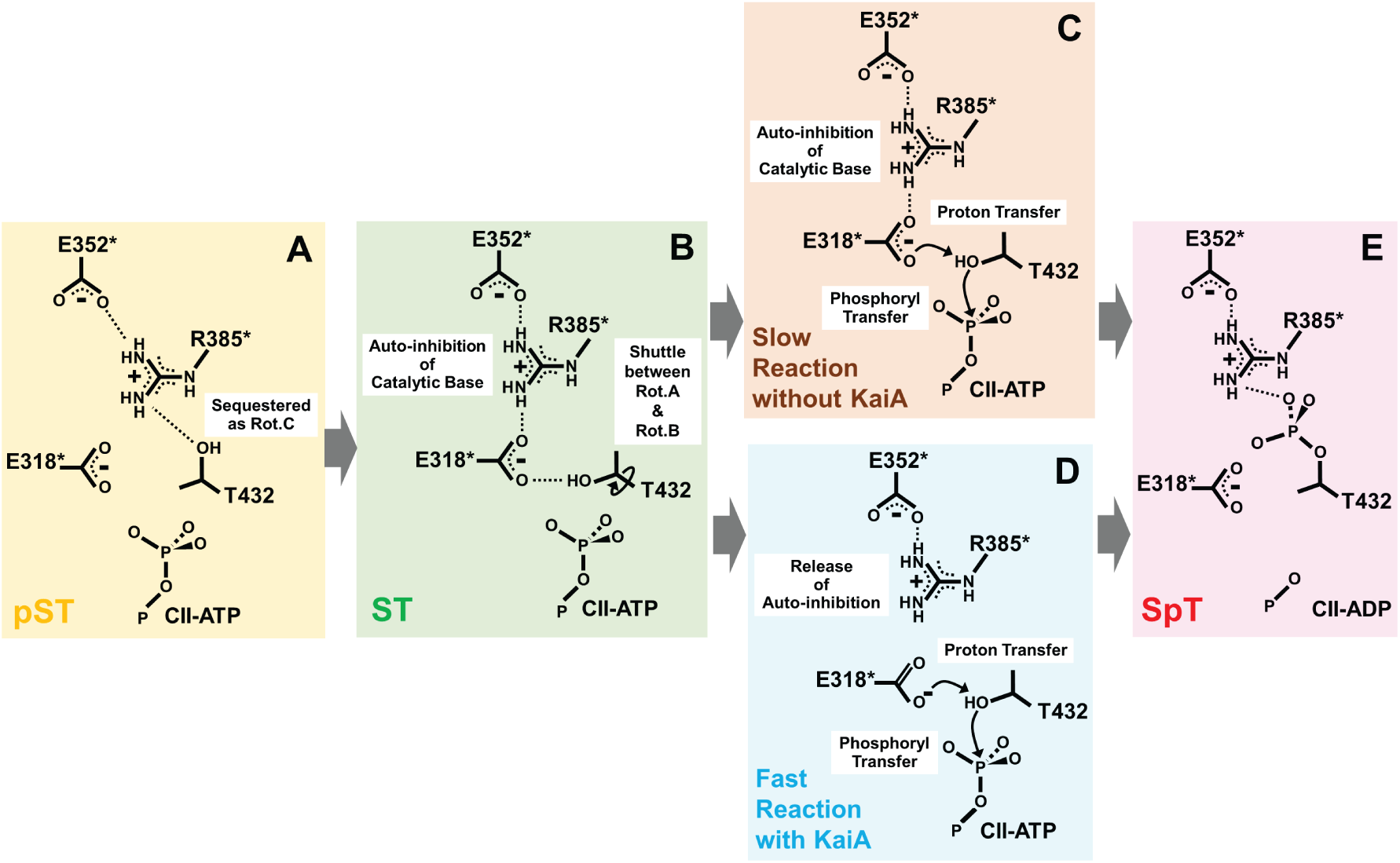
Proposed reaction scheme for priming phosphorylation at T432 in KaiC. (*A*) KaiC-pST, (*B*) KaiC-ST, (*C*) A reaction intermediate without KaiA, (*D*) the KaiA-assisted state, and (*E*) KaiC-SpT, respectively.

## Materials and Methods

### Expression and Purification of KaiC and its Mutants

Glutathione S-transferase (GST)-tagged KaiC was constructed in the plasmid vector, pGEX-6P-1. GST-tagged KaiC was expressed in *E. coli.* BL21(DE3) and purified as previously reported (14). The purified KaiC was dissolved in a buffer containing 20 mM Tris-HCl (pH8), 150 mM NaCl, 1 mM ATP, 5 mM MgCl_2_, 0.5 mM ethylenediaminetetraacetic acid (EDTA), and 1 mM dithiothreitol (DTT) (buffer A).

### KaiC Autodephosphorylation Assay

WT-KaiC and KaiC^R385A^ (0.6 mg/mL) was incubated at 30°C to induce autodephosphorylation. Aliquots were taken from the incubated samples at pseudo logarithmic intervals using the automated sampling device (54) to cover fast and slow relaxations quantitatively. The relative abundance of four phosphorylated KaiC forms was quantified by SDS polyacrylamide gel electrophoresis (SDS-PAGE) using LOUPE software (54). Kinetic traces for autodephosphorylation dynamics were fitted using a reversible four-state cyclic model as shown in Fig. 1*C* using MATLAB software.

### KaiC Autophosphorylation Assay

WT-KaiC was pre-incubated in buffer A for 50 h at 30°C to induce autodephosphorylation. The pH of the sample solution was changed using a buffer consisting of 20 mM Tris-HCl (pH9), 150 mM NaCl, 1 mM ATP, 5 mM MgCl_2_, 0.5 mM EDTA, and 1 mM DTT in a desalting column, PD MiniTrap G-25 (Cytiva). The sample solution at pH 9 (0.4 mg/mL) with abundant KaiC-ST was incubated at 30°C. Sampling and analyses were performed as described for the autodephosphorylation assay.

### KaiA-assisted Phosphorylation Assay

WT-KaiC and the KaiC mutants KaiC^R385A^, KaiC^E352Q^ and KaiC^E352D^ were pre-incubated in buffer A for 48 h at 30°C to induce autodephosphorylation. The KaiC solutions (0.2 mg/mL) were then mixed with KaiA (0.04 mg/mL) and the phosphorylation status was analyzed by SDS-PAGE.

### Analysis of the Crystal Structure of KaiC

In this study, the seven coordinates (PDB code: 7DYJ, 7DYE, 7DYI, 7DY1, 7DY2, 7DXQ, and 7V3X) deposited in the PDB databank were used for the analysis. The graphical presentations were generated by using Pymol.

### QM/MM and MD Simulations

The initial coordinates of the KaiC-ST hexamer with T432^rot-A^ were obtained from PDB: 7DYJ, and the loops that were missing from the crystal structure were modeled according to the previously reported structure (PDB: 2GBL) (17). The flexible N- and C-termini of KaiC-ST were omitted, i.e., residues I18–I497 of each KaiC-ST monomer were used. In addition, 12 ATP molecules, 12 magnesium ions, and crystal water molecules within the protein molecules were included in the system. Hydrogen atoms were added using the LEAP module in AmberTools. The KaiC-ST hexamer was surrounded by a water layer of 11.0 Å thickness and placed in a rectangular box. Sodium ions were added to neutralize the system charge. The total number of atoms in the system was thus approximately 207,000.

The Amberff99SB-ILDN force field (55) was used for the protein molecules, with modified parameters for ATP (56). The TIP3P model (57) was used for the water molecules. Long-range electrostatic interactions were treated using the particle mesh Ewald method (58), whereas short-range non-bonded interactions were cut off at 10 Å. A time step of 2 fs was used, and the simulations were performed using the Amber 2018 software (59) and the PLUMED 2.5.3 package (60).

The energy of each system was minimized for 1,000 steps while allowing only hydrogens to relax, followed by energy minimizations of 10,000 steps with restraints on backbone atoms to relax the sidechains and 10,000 steps without restraints. The temperature of the system was then gradually increased to 300 K over 100 ps while restraining the backbone atoms with a weak force constant of 1.0 kcal mol^-1^. Next, a constant-NPT (300 K, 1 atm) simulation was performed for 500 ps under the same restraint, followed by another 500 ps of constant-NPT (300 K, 1 atm) simulation without restraints. Finally, a 10-ns constant-NVT (300K) simulation was performed to complete the equilibration steps. The temperature was maintained using the Langevin thermostat with a collision frequency of 1.0 ps^-1^.

The mechanism of the phosphoryl transfer reaction was studied using the structure obtained after the equilibration steps described above. The interface between the protomers B and C, where the T432^rot-A^ O_γ_ – CII-ATP P_γ_ distance was kept short during the 1-µs trajectory, was used. The QM/MM method was used to study the chemical reaction. The QM region consists of the side chains of T432, E318*, and R385*, triphosphate and part of the ribose group (C5’, H5’1, and H5’2) of CII-ATP, and Mg^2+^. Link atoms are introduced to saturate the valence of the QM boundary atoms. To explore the conformational space efficiently, the density-functional-tight-binding method with third-order correction (DFTB3) (61) was chosen as the QM method. The timestep was decreased to 0.5 fs in the QM/MM simulations, and the system was equilibrated using the QM/MM Hamiltonian for 100 ps prior to the following calculations.

To obtain the initial reaction path starting from the reactant state, we performed a metadynamics (metaD) simulation (62) using the defined QM region. The difference (*q*_1_ = *r*_2_ – *r*_1_) between two distances, *r*_1_: T432^rot-A^ O_γ_ – CII-ATP P_γ_ and *r*_2_: CII-ATP P_γ_ – CII-ATP P_β_, was chosen as the reaction coordinate (Fig. 3*A*). Although we were interested in the breaking of the CII-ATP P_γ_ – CII-ATP O_3β_ bond, *r*_2_ was used to avoid the effect of permutations between the three CII-ATP O_β_ atoms. In the metaD simulation, a Gaussian potential of width = 0.20 Å and height = 0.10 kcal mol^-1^ was added every 0.1 ps. Half-harmonic potentials at *q*_1_ ≥ 5.7 Å and *q*_1_ ≤ 3.3 Å with a force constant of 100.0 kcal mol^-1^ Å^2^ were applied. The metaD simulation was performed for 300 ps.

To calculate the free energy profile of the reaction, replica exchange umbrella sampling (REUS) simulations were performed (63). The reaction was divided into two parts: the initial CII-ATP P_γ_ – CII-ATP O_3β_ bond dissociation, which leads to an intermediate state, and the subsequent T432^rot-A^ O_γ_ – CII-ATP P_γ_ bond formation coupled with the proton transfer from T432^rot-A^ O_γ_ to E318* O_ε_. In the first part, one-dimensional (1D) REUS was performed using *q_1_*, whereas in the second part, we performed two-dimensional (2D) REUS using two collective variables (Fig. 3*A*): *q*_1_ and the proton transfer variable *q*_2_ = *r*_4_ – *r*_3_, defined as the difference between two distances, *r*_3_: E318* O_ε_ – T432^rot-A^ H_γ_ and *r*_4_: T432^rot-A^ O_γ_ – T432^rot-A^ H_γ_. Harmonic restraints were applied along each coordinate with a force constant of 100.0 kcal mol^-1^ Å^-2^ in 1D-REUS, and 50.0 kcal mol^-1^ Å^-2^ and 25.0 kcal mol^-1^ Å^-2^ along *q_1_* and *q_2_*, respectively, in 2D-REUS. The restraints were centered at *q*_1_ = –1.0, –0.8,…2 in 1D-REUS, and *q*_1_ = 1.6, 1.8,…3.4 and *q*_2_ = –1.5, –1.0,…2.0 in 2D-REUS.

The number of replicas in the 1D- and 2D-REUS simulations were thus 16 and 80, respectively. The rate of exchange attempts between replicas was fixed at 1 ps in all REUS calculations. The initial structures of the replicas were taken from the snapshots along the metaD trajectory. In the 2D-REUS simulations, the initial structures were further generated by consecutively performing short (20 ps) 2D-umbrella sampling simulations from the metaD snapshots along the positive and negative *q*_2_-directions.

The replica was run for 820 and 620 ps in the 1D- and 2D-REUS simulations, respectively. The initial 120 ps was considered as the equilibration step and thus discarded from the following analyses. The 1D and 2D free energy surfaces (Fig. 3*C* and 3*D*) were constructed by the weighted histogram analysis method (64). The 1D free energy profile for the second part of the reaction was obtained as the minimum free energy path in the 2D free energy surface of *q*_1_ and *q*_2_.

To complement the QM/MM simulations and to explore the conformational states, e.g., the T432 rotamers in solution, MD simulation using classical mechanical force fields was performed from the equilibrated structure described above. The time step was set to 2 fs, and the MD simulation was performed under the constant-NVT (300K) condition for 1 µs.

## Acknowledgments

The calculations are partially carried out at the Research Center for Computational Sciences in Okazaki (23-IMS-C111 to T.M. and 23-IMS-C196 to S.S.).

## Data Availability

All data relating to this study have been presented in the manuscript and supplementary materials.

