## Supplementary material for "The priming phosphorylation of KaiC is activated by the release of its autokinase autoinhibition": SI Appendix

**This PDF file includes:**

Figures S1 to S7
Tables S1

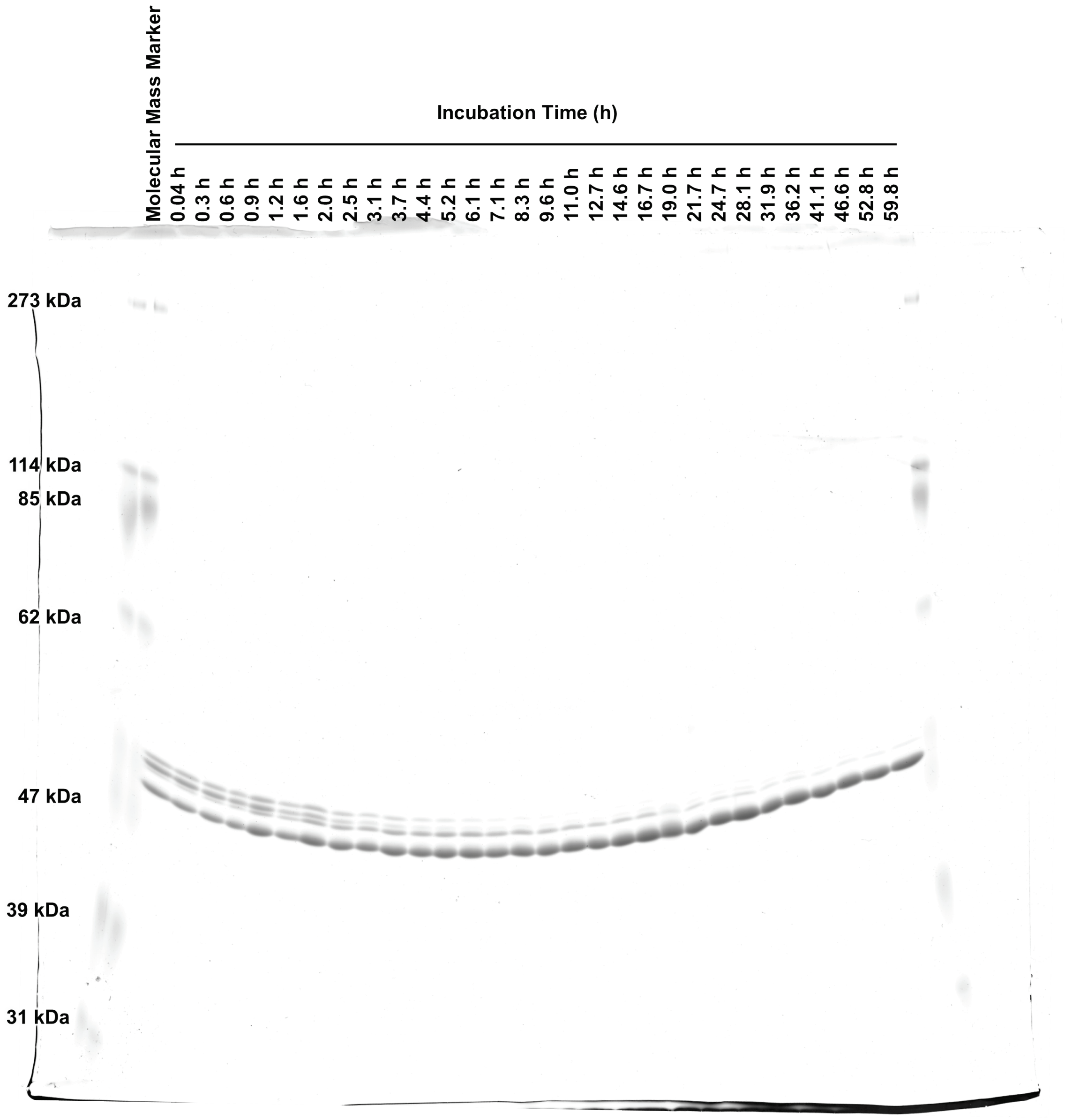


**Fig. S1.** Image of the original SDS-PAGE gel displayed in Fig. 1*A*.

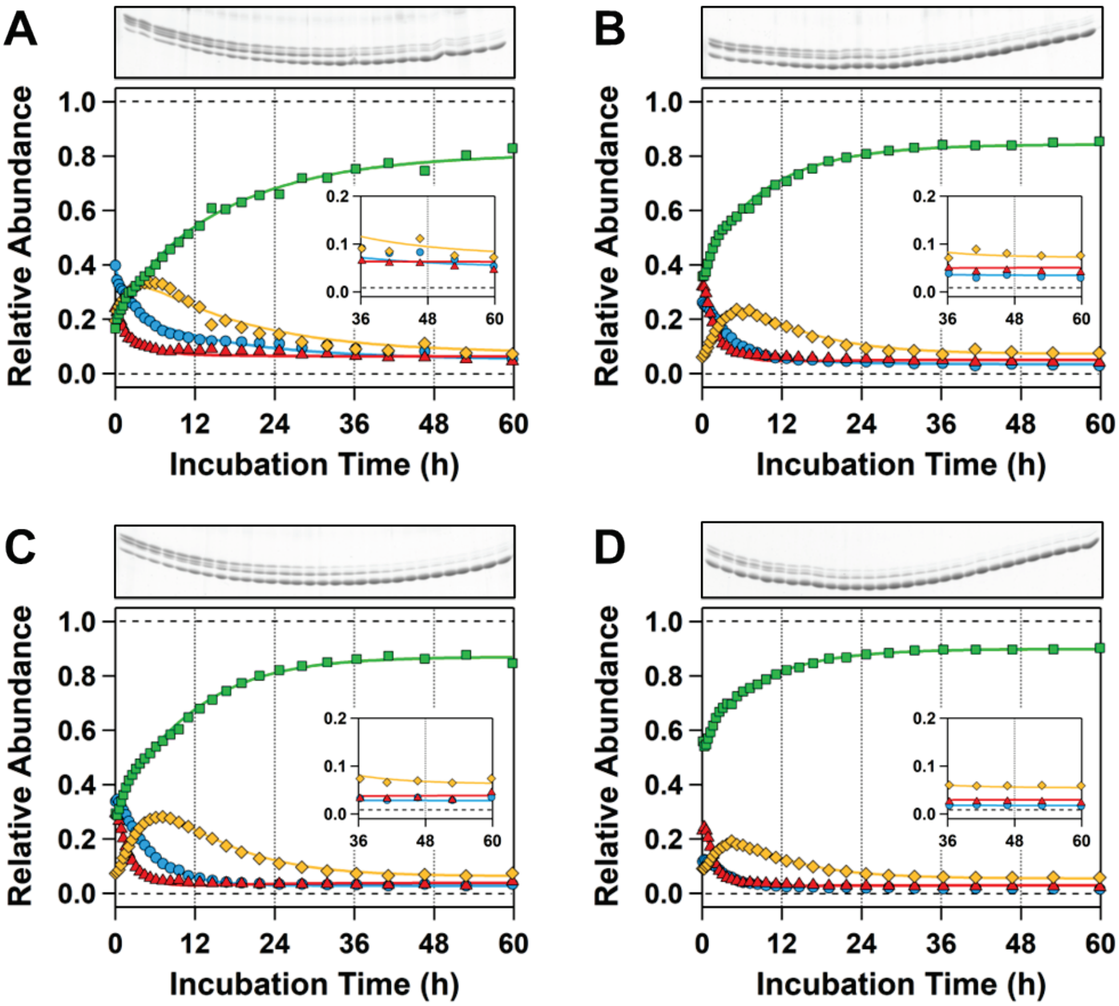


**Fig. S2.** Autodephosphorylation dynamics. (*A*–*D*) The reaction was initiated with four different relative abundances of the ST, SpT, pSpT, and pST forms of KaiC. Solid lines indicate the best fit from global nonlinear least-squares analysis using the four-state model shown in Fig. 1*C*. The insets show the enlarged plots and fits of the region corresponding to the longer periods. Converged rate constants are compiled in Table S1.

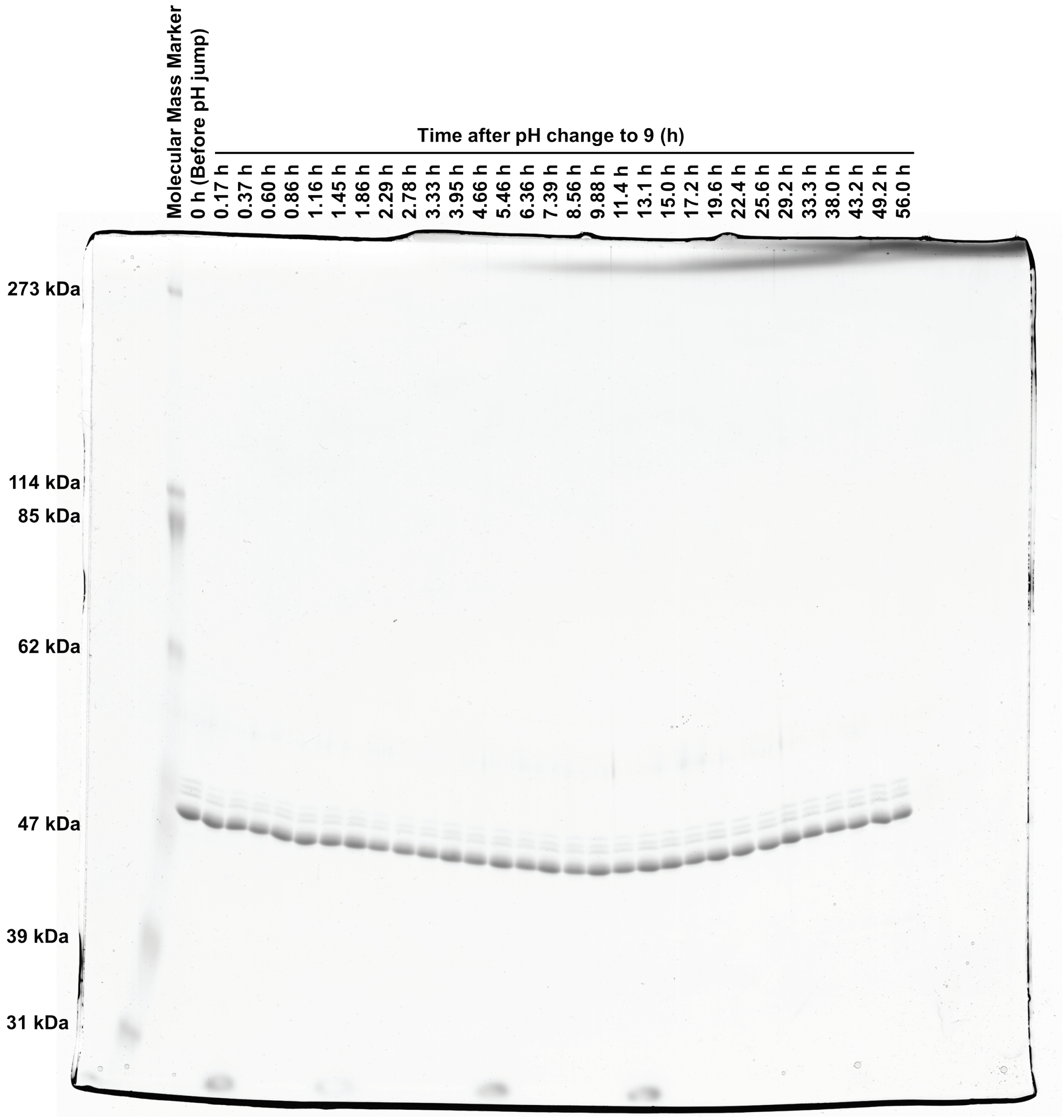


**Fig. S3.** Image of the original SDS-PAGE gel loaded with samples from KaiC under autophosphorylation conditions at 30°C after increasing the pH from 8.0 to 9.0.

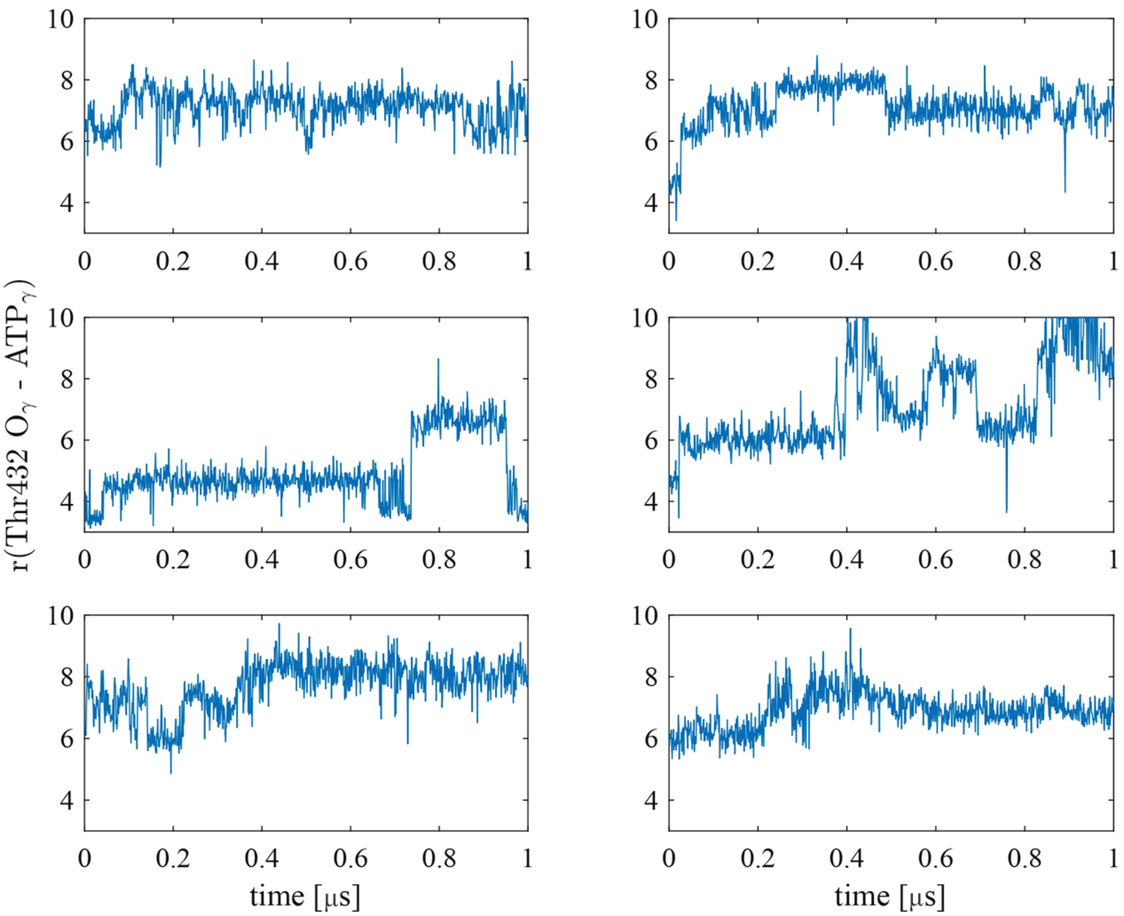


**Fig. S4.** Distances between CII-ATP $P_\gamma$ and T432 $O_\gamma$ during the 1 μs MD trajectory. The CII-ATP $P_\gamma$ – T432 $O_\gamma$ distance of approximately 5 Å or less corresponds to the rotamer A of the T432 side chain (middle left panel).

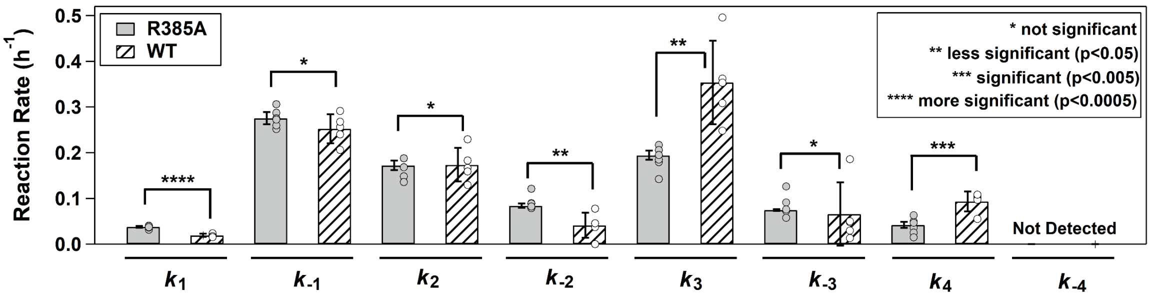


**Fig. S5.** Effects of the R385A substitution in KaiC-ST on the eight rate constants shown in Fig. 1*C*.

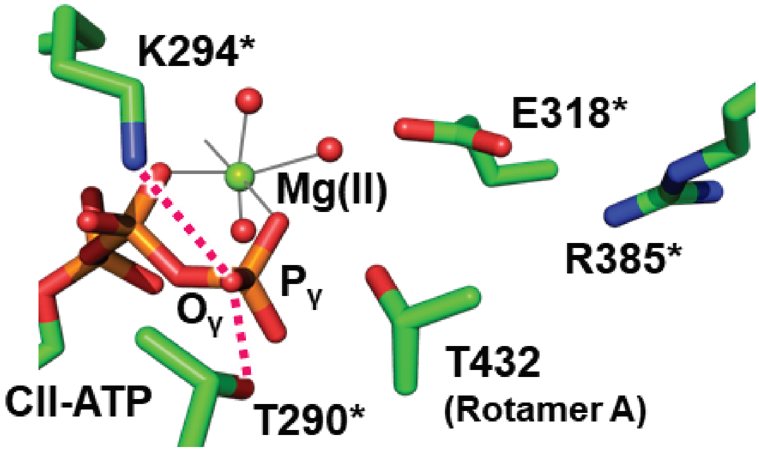


**Fig. S6.** Zoomed-in view of CII-ATP $O_\gamma$ stabilized by hydrogen bonding with T290* and electrostatic interaction with K294* in KaiC-ST (PDB: 7DYJ).

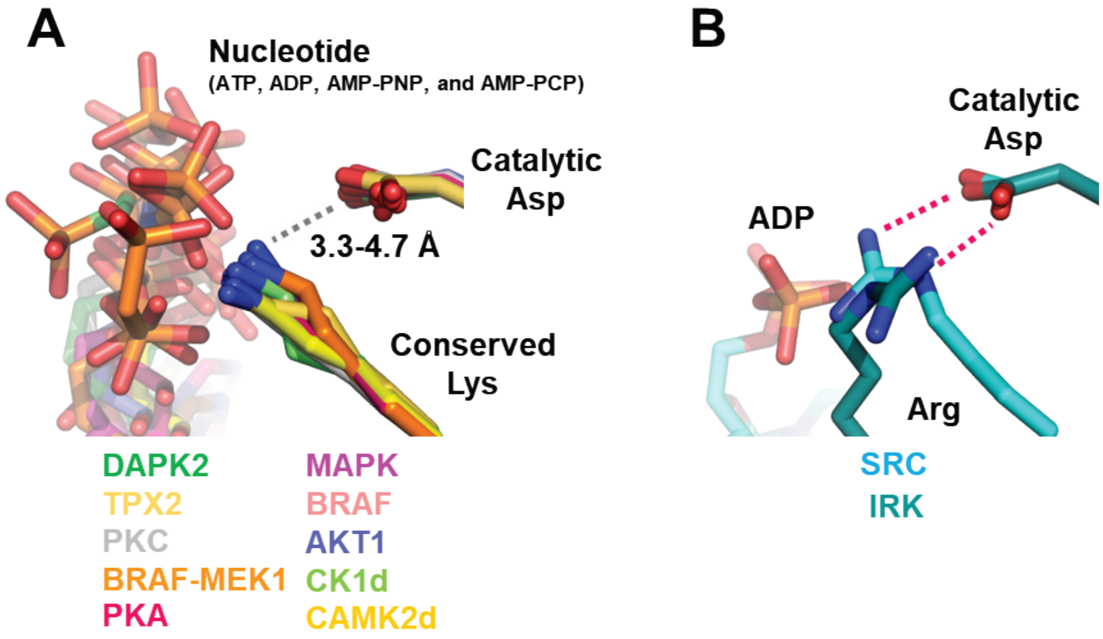


**Fig. S7.** Zoomed-in view of the active sites of representative protein kinases. (*A*) The nucleotide-bound forms of DAPK2 (PDB code: 2YAA), TPX2 (PDB code: 1OL5), PKC (PDB code: 3A8W), BRAF-MEK1 complex (PDB code: 4MNE), PKA (PDB code: 2QCS), MAPK (PDB code: 6NYB), BRAF (PDB code: 1UWH), AKT1 (PDB code: 4EKK), CK1d (PDB code: 6RU6), and CAMK2d (PDB code: 2VN9) are shown as examples in which each catalytic Asp is not involved in a stable hydrogen bond with a positively charged Lys nearby. (*B*) ADP-bound form of SRC (PDB code: 6F3F) and nucleotide-free form of IRK (PDB code: 1IRK) are shown as examples in which each catalytic Asp is hydrogen bonded with a positively charged Arg.

**Table S1.** Rate constants determined by global nonlinear least-squares analysis using the four-state model shown in Fig. 1*C*.

| | Rate Constants ($h^{-1}$) | | | | | | | |
|---|---|---|---|---|---|---|---|---|
| | $k_1$ | $k_{-1}$ | $k_2$ | $k_{-2}$ | $k_3$ | $k_{-3}$ | $k_4$ | $k_{-4}$ |
| **Fig. 1*B*** | 0.019 | 0.291 | 0.183 | 0.045 | 0.362 | 0.030 | 0.105 | 0 |
| **Fig. S2*A*** | 0.021 | 0.206 | 0.130 | 0.078 | 0.353 | 0.186 | 0.055 | 0 |
| **Fig. S2*B*** | 0.024 | 0.256 | 0.167 | 0.037 | 0.309 | 0.048 | 0.099 | 0 |
| **Fig. S2*C*** | 0.018 | 0.240 | 0.159 | 0 | 0.248 | 0.013 | 0.097 | 0 |
| **Fig. S2*D*** | 0.015 | 0.268 | 0.229 | 0.045 | 0.496 | 0.050 | 0.109 | 0 |
| **Average** | 0.019 | 0.252 | 0.173 | 0.041 | 0.354 | 0.065 | 0.093 | 0 |
| **S.D.** | 0.003 | 0.032 | 0.037 | 0.028 | 0.092 | 0.069 | 0.022 | 0 |
